# Glycoconjugate diversification in *Campylobacter concisus* is determined by two glycosyltransferases

**DOI:** 10.64898/2026.05.23.727390

**Authors:** Hayley L. Knox, Christine A. Arbour, Chaoshuang Xia, Catherine E. Costello, Barbara Imperiali, Karen N. Allen

**Affiliations:** Department of Chemistry, Boston University, Boston, MA 02215 USA; Department of Biology, Massachusetts Institute of Technology, Cambridge, MA 02139 USA; Department of Chemistry, Massachusetts Institute of Technology, Cambridge, MA 02139 USA; Center for Biomedical Mass Spectrometry, Boston University Chobanian & Avedisian School of Medicine, Boston, MA 02118 USA

## Abstract

Bacterial glycoconjugates are structurally diverse, with enormous variation in sugar identity, modifications and linkages. Glycoconjugates play key roles in numerous cell functions, acting as the primary interface with the environment. Asparagine (*N*)-linked glycosylation has been extensively studied in the pathogenic *Campylobacter* genus, due to the availability of numerous genome sequences and the highly conserved pathway logic, despite the final *N*-linked glycan product diversity. We recently reported on a partitioning of *N*-linked glycan structures between the *Campylobacter* species, focused on the inclusion of a C6-carboxyl-sugar in the third position of the growing glycan in *Campylobacter concisus*. However, at the time, the final glycan was not fully defined. Here, we identify the final glycan product in *C. concisus*, demonstrating surprising substrate promiscuity of the GT-A enzyme, PglI, which adds the penultimate sugar, and uncover a previously uncharacterized enzyme (GT-25) that unexpectedly adds the final sugar to complete the heptasaccharide product. Through a detailed study of these two *C. concisus* pathway enzymes, the intermediate and final glycans were defined, with determination of linkage positions of major and minor isomeric products following each glycan addition, through high-resolution electronic excitation dissociation tandem mass spectrometry. These findings on the Group II *C. concisus N*-linked glycan highlight the diversification of the glycan and the utilization, at the non-reducing end, of GlcNAc over GalNAc, which is dominant in the Group I species.

## Introduction

Bacterial glycans are remarkably diverse biomolecules, exceeding the complexity observed in eukaryotes (1). In bacteria, there is a broad repertoire of sugars available, including sugars specific to prokaryotes. This diversity is further expanded upon with variable linkage stereochemistry, branching, and chemical modifications (2). These glycans are assembled into essential cellular components including lipopolysaccharides, capsular polysaccharides, peptidoglycans, and *N*-and *O*-linked glycoproteins, amongst others (3). Notably, the glycans play a crucial role in the cell-surface structures which are vital to the survival of bacteria and for mediation of crucial interactions with other bacteria and the hosts that they infect (4,5). Because bacterial glycoconjugates provide the primary interface with the environment (including antibiotics), increased structural diversity of the sugars through sugar modification and glycoform variation can be crucial for bacterial survival and virulence (1,6,7).

In bacteria, *N*-linked glycosylation at selected asparagine residues in proteins is responsible for mediating numerous functions, including cell-cell interactions, colonization, and pathogen-host interactions (4,5). Additionally, the cell-surface *N*-linked glycans play a key role in bacterial virulence and pathogenicity (8,9). Intriguingly, the *N*-linked glycoconjugate biosynthetic pathway is largely conserved across the *Campylobacter* genus, as evidenced by the availability of more than 40 complete genome sequences (10), although there is diversity in the final distribution of glycoforms. Members of the *Campylobacter* genus, particularly *Campylobacter jejuni* (*Cj*), are amongst the leading global causes of bacterial gastroenteritis in humans and animals (11–13). The importance of the *N*-linked glycoconjugate biosynthetic pathway in *Campylobacter* is emphasized by reduced levels of bacterial colonization in chicken hosts upon disruption of the glycan assembly pathway (8). The *N*-linked protein glycosylation (*pgl*) pathway is initiated by a phosphoglycosyltransferase (PGT) that transfers a soluble phosphosugar to a membrane-anchored polyprenyl-phosphate (undecaprenyl) acceptor. The polyisoprenyldiphosphate-linked glycan is elongated and elaborated by glycosyltransferases (GTs) that act sequentially. The GTs (either GT-A or GT-B fold) catalyze the stereospecific formation of glycan linkages. The final glycan transfer across the membrane is catalyzed by a flippase (PglK) prior to the glycan being appended to the appropriate asparagine by an oligosaccharide transferase, designated as PglB.

The *pgl* pathway has been extensively characterized in *Cj* and comprises an 11-gene operon (**Figure 1A**) (5,14–18). In that pathway, the final UndPP-linked glycoconjugate is GalNAc-α1→4-GalNAc-α1→4(Glc-β1,3)-GalNAc-α1→4-GalNAc-α1→4-GalNAc-α1→3-diNAcBac-α-1-diphosphate undecaprenyl (PP-Und) (**Figure 1B**) (14,19–22). Notably, the assignment of anomeric configurations was achieved by NMR (21). Closely related *Campylobacter* species have a similar operon organization, but the precise function of the gene products vary (**Figure 1A**). In previous studies on the *Campylobacter concisus* (*Cc*) pathway, the final glycan product was not conclusively identified. From an *in vitro*-generated glycan study utilizing biochemical analysis and mass spectrometry, it had been concluded that the final glycan is identical to that from *Cj*, with the exception of a sugar with a mass of 217 Da replacing GalNAc at the position catalyzed by PglJ (20). Previous work from our laboratories focused on characterizing the glycosyltransferases PglC through PglH1 from *Cc*, which catalyze the addition of the first five carbohydrates. The glycan produced by those enzymes was identified as GlcNAc-GlcNAc-GalNAcA-GalNAc-diNAcBac-PP-Und, with anomeric linkages assigned as analogous to those in *Cj,* based on glycosyltransferase fold and sequence identity (23,24). Though the corresponding enzymes in the *Cj* and *Cc* pathways are similar in sequence (∼40% identity), there are significant differences in the final glycan structures (**Figure 1C**). The enzymes catalyzing the initiating PGT reaction and the first glycosyltransferase reaction, PglC and PglA are orthologs, appending the same sugars in *Cj* and *Cc* (24). The divergence in the two pathways begins with PglJ. In *Cj*, PglJ appends GalNAc whereas in *Cc*, a sugar with a mass of 217 Da, later identified as the C6-oxidized sugar, GalNAcA was transferred to the growing glycan (24). This finding was consistent with the proposed split of group I and group II strains initially proposed during a comparative genomic study of the original 30 sequenced *Campylobacter* taxa, which grouped the bacteria based on similarity to *Cj* (group I) and more diverse final glycans (group II) (19). The *pgl* pathway becomes more complicated after the action of PglJ. In *Cj*, a single enzyme, PglH, appends three GalNAc residues consecutively to the growing glycan chain (14,19). *Cc* encodes two enzymes with similarity to PglH, PglH1 and PglH2 (23). Through kinetic and bioinformatic analyses, it was determined that each enzyme appended a single GlcNAc, with PglH2 acting first, followed by PglH1 (23). It was hypothesized that the inclusion of the GalNAcA sugar, transferred by PglJ, forced the divergence of the PglH enzymes by changing the sugar acceptor (23). Overall, these findings confirmed the identity of the sugars appended by PglJ, PglH2, and PglH1 in *Cc*, however, the final two HexNAc additions remained to be characterized.

**Figure 1.**
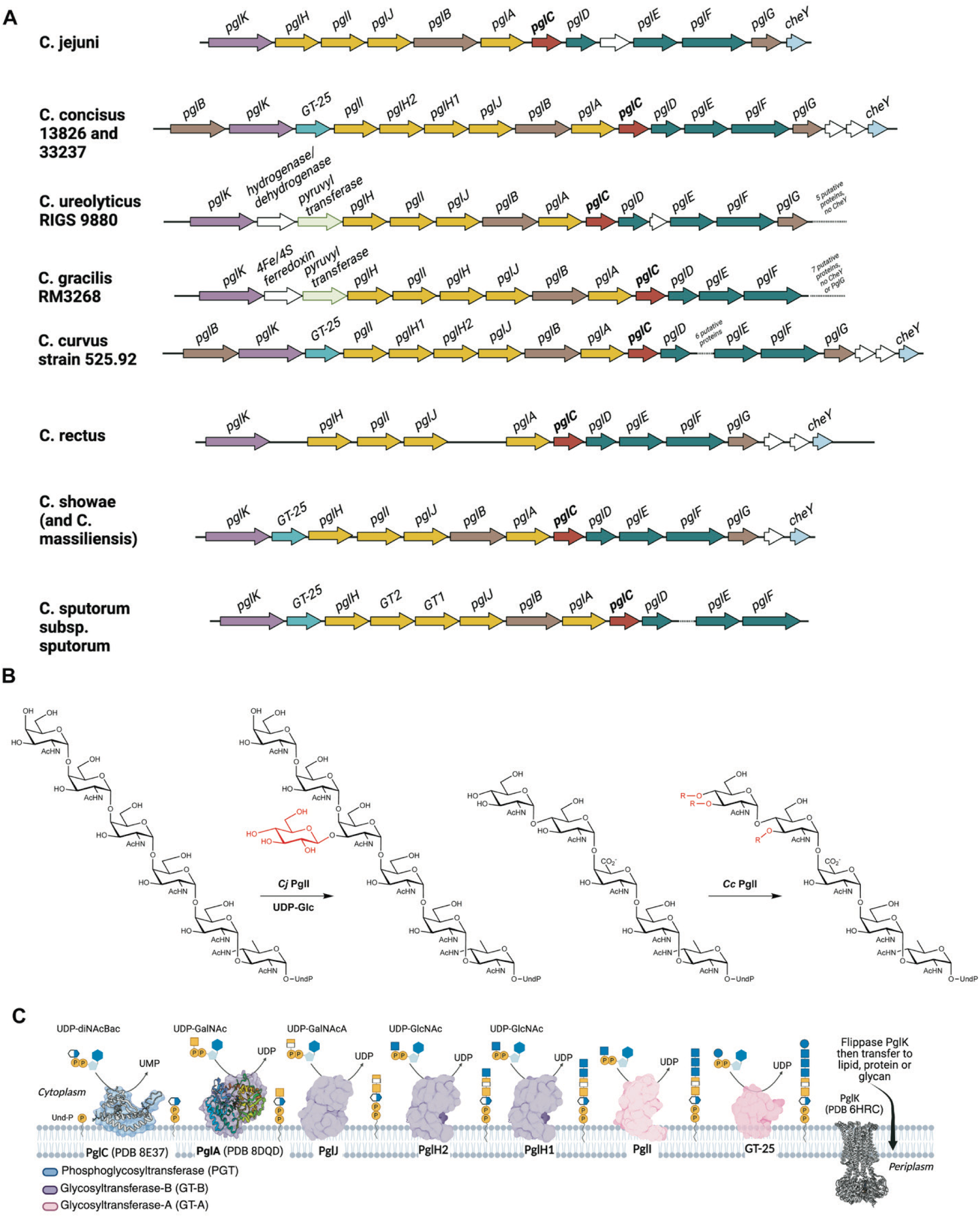
The *pgl* pathway. (**A**) Genome neighborhood diagram of a representative group of *Campylobacter* species. (**B**) Reaction scheme of the glycosyltransfer reaction carried out by *Cj* PglI and *Cc* PglI. (**C**) Schematic of the *pgl* pathway in *Cc*.

In *Cj*, the GT-A fold enzyme PglI is the final GT in the pathway, appending a β-1→3Glc to the hexasaccharide (14) (**Figure 1B**). In *Cc*, there is an enzyme annotated as PglI that shares ∼30% identity with PglI from *Cj* and includes the same hallmark sequence motifs. Intriguingly, within the *Cc pgl* operon, a gene encoding a glycosyltransferase family 25 (*gt-25*) enzyme is positioned between *pglK* and *pglI* (**Figure 1A**). The GT-25 enzyme also exhibits a GT-A fold, but has no confirmed role in the *pgl* pathway. GT-25 shares low sequence identity with PglI (22.7%); although the Rossmann-like domains that defines the GT-A family do align, including the canonical GT-A family active site motif (DXD), the C-terminal domains do not. Notably, in *Campylobacter*, the *gt-25* gene is often found in operons where the glycan biosynthetic pathway has diverged to include C6-carboxyl hexosamine sugars in the final product (**Figure 1A**). Herein, the specific biochemical functions of *Cc* PglI and GT-25 were determined through bioinformatic analysis, biophysical, and biochemical assays (**Figure 1C**). The glycan structures, including connectivity and topology were determined by high-resolution electronic excitation dissociation (EED) tandem mass spectrometry. This allowed for the discrete identification of the distinct structure(s) of the products following each glycan addition and revealed how PglI and GT-25 each add diversity to the final glycoconjugate through their substrate promiscuity.

## Materials and Methods

### Expression and purification of PglC, PglA, PglJ, PglH2, PglH1, PglH1 S258P/N259T variant, PglI, and GT-25

PglC, PglA, PglJ, PglH2, PglH1, and the PglH1 *S258P/N259T* (SN_PT) variant were expressed and purified as previously described (23–25). It is important to note that the PglI orthologs from *C. concisus* strain 13826 and strain 33237 are 96% identical and the GT-25 orthologs are 94% identical and thus are treated as equivalent. PglI from the *C. concisus* strain 33237 (uniprot ID: A0A0M4SNM0) and glycosyltransferase family 25 enzyme (GT-25) (uniprot ID: A7ZEU1) from *C. concisus* strain 13826 were produced heterologously in *E. coli* BL21 (DE3) by overexpression from a pET28a:*pglI* or pET28a:*gt-25* construct using the same purification protocol. The genes were codon-optimized, synthesized, and cloned by Twist Bioscience (San Francisco, USA). Cells were cultured in autoinduction media supplemented with 150 μg/mL kanamycin at 37 °C at 225 rpm for 20 hours. Once the cultures reached OD_600_ ∼0.8, they were incubated at 18 °C for 16 hours before the cells were collected by centrifugation at 5000*g*. The resulting cell paste (approximately 20 g per liter of culture) was stored at -80 °C prior to purification.

For protein isolation, the cell paste was resuspended in 50 mL lysis buffer (50 mM HEPES pH 7.5 and 300 mM NaCl). The cell suspension was incubated with lysozyme (1 mg/mL), DNase (0.05 mg/mL), and EDTA-free protease inhibitor cocktail set III (1 µl/mL, Millipore Sigma). The cells were lysed by microfluidization (at 18,000 PSI) and centrifuged at 6926*g* for 45 min to remove cell debris. The supernatant was transferred to clean ultracentrifugation tubes and centrifuged at 104,361*g* for 65 min to sediment the cell envelope fraction (CEF). The CEF was homogenized using a Dounce tissue homogenizer in ∼20 mL lysis buffer. The concentration of the resuspended CEF was checked by OD_280_ and the final volume was adjusted such that the CEF protein concentration was 50 mg/mL. The CEF preparation was flash frozen in liquid nitrogen and either purified immediately or, stored at -80 °C.

The detergent *n*-dodecyl-β-D-maltoside (DDM) was added to the CEF preparation to a final concentration of 1% to solubilize the enzyme of interest. The solution was rotated for approximately 1 hour at 4 °C. The solution was then clarified by centrifugation at 141,261*g* for 65 min to afford the detergent-soluble fraction. This fraction was loaded onto a pre-equilibrated column of Ni-NTA resin (2 mL) and purified by immobilized-metal affinity chromatography. The protein-loaded resin was washed with 4 column volumes (CVs) of wash 1 buffer (lysis buffer supplemented with 25 mM imidazole, 5% glycerol, and 0.03% DDM) and 4 CVs of wash 2 buffer (lysis buffer supplemented with 50 mM imidazole, 5% glycerol, and 0.03% DDM) before the addition of 5 CVs of elution buffer (lysis buffer supplemented with 500 mM imidazole, 5% glycerol, and 0.03% DDM). Fractions containing enzyme were pooled and concentrated to 2.5 mL in an Amicon 10 kDa MWCO ultrafiltration device (Millipore Sigma). The protein fractions were exchanged into gel filtration buffer (25 mM HEPES pH 7.5, 150 mM NaCl, 5% glycerol, and 0.03% DDM) using a PD-10 pre-poured gel-filtration column (Cytiva) and concentrated prior to flash cooling in liquid nitrogen for storage at -80 °C.

### Expression and purification of GT-25 AAA variant

The GT-25 AAA variant (E^90^D^91^D^92^ to A^90^A^91^A^92^) was synthesized as a gene fragment (Twist Bioscience) with overhangs for insertion into a vector used for a previously described purification strategy (26). The plasmid includes a SUMO tag (to aid with solubility) and a dual-Strep Tag for purification. The GT-25 AAA variant was transformed into competent C43 cells containing pAM174, encoding an arabinose-inducible Ulp1 protease for cleavage of the SUMO tag *in vivo*. Expression was carried out as described for PglI and GT-25, with the addition of 25 µg/mL of chloramphenicol (GoldBio) included in the media. At OD_600_ ∼0.8, 1 g of arabinose (GoldBio) was added prior to lowering the temperature.

Expression and extraction of the GT-25 AAA variant was carried out as for the wild-type enzyme. For purification, 1 mL of Streptactin-XT resin (IBA Lifesciences) was equilibrated with 15 mL of lysis buffer, and the post-extraction supernatant was run over the column bed three times. The column was washed with 10 mL of lysis buffer and then 6 mL of strep elution buffer (50 mM HEPES pH 7.5, 300 mM NaCl, 5% glycerol, 50 mM biotin, 0.03% DDM) was incubated on the resin for >30 min prior to elution. The protein eluate was concentrated to ∼2.5 mL with an Amicon filter (Millipore Sigma) concentrator, then exchanged over a PD-10 column in gel filtration buffer and concentrated prior to being flash frozen in liquid nitrogen for storage at -80 °C.

### Chemoenzymatic synthesis reactions

Various reactions were set up (see **Table S1**) for chemoenzymatic synthesis. Purified enzymes were added at an approximate concentration of 1 µM, apart from PglI and GT-25, which were added at approximate concentrations of 4 µM and 5 µM, respectively. UndP was used at a 200 µM concentration from a 20 mM stock in DMSO; the final concentration of DMSO in each reaction was 10%. The following nucleotide sugars were added, when appropriate, at the following concentrations: 300 µM UDP-diNAcBac, 300 µM UDP-GalNAc, 300 µM UDP-GalNAcA, 1 mM UDP-GlcNAc or UDP-GlcNAc[^2^H]_3_, and 600 µM UDP-Glc. Reactions typically had a final volume of 200 µL.

Reactions were allowed to proceed for either 2 hours or overnight. Reactions were quenched with 2 mL of CHCl_3_/MeOH (2:1). The separated lower organic layer was washed three times with 500 µL of either PSUP (pure solvent upper phase, 15 mL CHCl_3_, 240 mL MeOH, 1.83 g KCl, and 235 mL of H_2_O) or salty PSUP (pure solvent upper phase, 240 mL MeOH, 16 g NaCl, 240 mL H_2_O), and then the organic layer was dried down under a N_2_ gas stream for further processing. PSUP was used for trisaccharides and shorter saccharides and salty PSUP was used for tetrasaccharides and larger saccharides.

### Fluorescence labeling and 2-aminobenzamide (2-AB) characterization

Reducing-end 2-AB labeling was performed as previously established (14,23,27). Briefly, the extracted glycan-PP-Und products were hydrolyzed with a 1:1 (v/v) mixture of *n*-propanol and 2 M trifluoroacetic acid at 50 °C for 15 min to generate glycans with a free reducing-end. This solution was evaporated to dryness. To generate the 2-AB labeling reagent, 5 mg of 2-aminobenzamide (Alfa Aesar, product #A14756) was dissolved in 100 µL of acetic acid/DMSO (1:2.3, v/v). This solution was used to dissolve 6 mg of sodium cyanoborohydride (Sigma-Aldrich, product #156159) to yield the 2-AB labeling reagent. Then 5 µL of the mixed 2-AB/sodium cyanoborohydride solution was used to resuspend the dried saccharides and the solution was incubated at 60 °C for 2-4 hours. The solution was diluted with 50 µL of H_2_O and purified by HPLC with fluorescence detection. An analytical reversed-phase HPLC column (Prozyme GlycoSepR, GK14727, currently sold by Agilent with part number NC9421472) with a 50 mM ammonium acetate (pH 4.4) system was used to separate the product from excess dye (solvent A: 50 mM ammonium acetate, 10% methanol; solvent B: 50 mM ammonium acetate, 20% methanol). A mobile phase gradient from 0-100% solvent B was run over 40 min at a flow rate of 0.7 mL/min. Peak elution was monitored with a fluorescence detector (Agilent Fluorescent Detector, G1321A) with λ_ex_ = 330 nm and λ_em_ = 420 nm. Peaks were collected, lyophilized, and analyzed by ESI-MS in the negative mode on an Agilent 6125B quadrupole mass spectrometer (detection range *m/z* 100 to 1500) attached to an Agilent 1260 Infinity LC prior to transfer to the Boston University School of Medicine Center for Biomedical Mass Spectrometry for detailed analysis.

### Enzyme assays

*Cc* PglI and GT-25 substrate specificity was evaluated using a radiolabeled UDP-[^3^H]-sugar panel and an extraction-based assay, as previously described (23,27,28). Luminescence-based assays for PglI and GT-25 were performed using the UDP-Glo assay (Promega) to detect UDP production. The quenching solution was prepared as described by Promega. The UDP-Glo standard curve was obtained at UDP concentrations of a serial dilution with starting concentration of 25 µM (25, 12.5, 6.25, 3.12, 1.56, 0.79 µM) from 10X UDP stocks and included 10% DMSO. Reactions included approximately 20 µM GlcNAc-GlcNAc-GalNAcA-GalNAc-diNAcBac-PP-Und acceptor (10% DMSO final), 50 mM HEPES pH 7.5, 100 mM NaCl, 5 mM MgCl_2_, 0.1% Triton X-100, and 1 µM of PglI, GT-25, or GT-25 AAA variant in various combinations. Reactions were initiated with the addition of 500 µM UDP-GlcNAc and allowed to proceed for 20 min prior to quenching with 11 µL of quenching solution. Each product mixture was transferred to a 96-well plate (white, non-binding surface, Corning). The plate was shaken at a low speed for 30 s and incubated for 1 hr at 25 °C. Luminescence was measured on a plate reader (BioTek Synergy H1 Hybrid Reader).

### Nanoscale Differential Scanning Fluorimetry (nanoDSF)

GT-25 was analyzed using nanoDSF (Prometheus Panta, Nanotemper) at a concentration of 1 mg/mL and 2 mM sugar donor concentration. Analysis was carried out in 25 mM HEPES pH 7.5, 150 mM NaCl, 5% glycerol, and 0.03% DDM. GT-25 was preincubated with the substrate on ice for 15 min prior to measurement. Protein was analyzed in duplicate using an excitation wavelength of 280 nm and applying 100% excitation power. A heating curve of 1.5 °C/min over a temperature range of 20 to 95 °C was used. The first derivative of the 350 nm/330 nm ratio was used for analysis.

### Pull-down assays of PglH1, PglI, and GT-25

Pull-down assays for PglH1, PglI, and GT-25 were conducted. For this study, pet26b-PglH1, dual Strep-tagged PglI, and pet28a-GT-25 were utilized. The CEF (0.5 mL) at 50 mg/mL of the appropriate enzyme combinations were mixed and incubated with styrene maleic acid (SMA) 300 for 1 hr while rotating. The mixed CEF was applied to 0.5 mL Streptactin-XT resin preequilibrated in 50 mM HEPES pH 7.5, 200 mM NaCl. The resin was washed with 6 CVs of the equilibration buffer. Sample was eluted in 2 CVs of equilibration buffer with 50 mM biotin added. The resulting fractions were run on SDS-PAGE after standard denaturation. The samples were: (1) PglH1+PglI, (2) PglH1 (control), (3) PglI+GT-25, and (4) GT-25 (control).

### Bioinformatic analyses

The SSNs for PglI and GT-25 were constructed using the EFI-Enzyme Similarity Tool (EFI-EST), available at https://efi.igb.illinois.edu/efi-est/ (29,30). For both, the query input (PglI: A0A0M4SNM0 and GT-25 A7ZEU1) was used for BLAST analysis, constructing an SSN of approximately 10,000 sequences. The Enzyme Function Initiative-Genome Neighborhood Tool (EFI-GNT, https://efi.igb.illinois.edu/efi-gnt/) was used to generate GNDs (Genome Neighborhood Diagrams), using the GT-25 SSN as input (30).

### High Resolution mass spectrometry (MS) and tandem MS (MS/MS) analyses

The 2-AB-labeled glycan samples were delivered through direct-infusion nanoESI in the positive mode via a TriVersa NanoMate (Advion, Ithaca, NY) and analyzed on a Fusion Lumos Tribrid Orbitrap MS (Thermo Scientific, Waltham, MA) to obtain initial MS and higher energy collisional dissociation (HCD) MS/MS spectra, to measure the accurate *m/z* values and verify the topology of the major components, and then on a NanoMate-equipped Q Exactive-HF orbitrap instrument (Thermo Scientific, Waltham, MA) modified with an Omnitrap (FasmaTech, Athens, Greece) to establish the positions of glycosidic linkages and distinguish among isomeric products. For the electronic excitation dissociation (EED) MS/MS experiments described herein, ions of interest were isolated in the QE-HF quadrupole and then irradiated with electrons in the Q5 trap of the Omnitrap before being sent back to the orbitrap for m/z detection (31). The electron energy at the Omnitrap was tuned between 17 and 20 eV and optimized for each experiment. The electron irradiation time was typically 100-400 ms for EED experiments. All spectra were acquired with 10 microscans and a maximum injection time of 800 ms.

### Data Analysis

Data were processed using the Xcalibur software (Version 4.1.50, Thermo Scientific, Waltham, MA). Manual data interpretation was assisted by ChemDraw software (Revvity Signals). Peaks were generally assigned with a mass accuracy within 3 ppm. Fragmentation annotation was based on the Domon and Costello nomenclature for the glycan motif (32).

## Results and discussion

### Characterization of Cc PglI and GT-25

Our previous work explored the presence and specificity, of two GT-B fold glycosyltransferases PglH1 and PglH2 in *Cc* and the majority of Group II species (23). In Group I species such as *Cj*, there is a single PglH GT and this enzyme is processive, adding three GlcNAc residues to the trisaccharide glycoconjugate (14). Although PglH1 and PglH2 are similar in sequence to *Cj* PglH (40% and 39% sequence similarity, respectively), the enzymes are not processive and each adds a single GlcNAc (23). Herein, we sought to determine the final glycan product of *Cc* and to understand the role of PglI in this pathway. We identified a new enzyme (GT-25) that unexpectedly plays a role in glycan synthesis.

The composition of the *Cc* glycan was originally characterized through tandem mass spectrometry and suggested the linear sequence of three repeating HexNAcs linked to the reducing-end trisaccharide (19,20). *In vitro* biochemical analysis with purified enzymes and substrates in our laboratories has confirmed the sugar preferences for the enzymes PglC, PglA, PglJ, PglH2, and PglH1 and proposed that the product generated from those enzymes is GlcNAc-GlcNAc-GalNAcA-GalNAc-diNAcBac-PP-Und (23,24). Because it appears that neither PglH2 nor PglH1 catalyzes addition of more than one GlcNAc residue, it was unclear which enzyme in the operon installs the third HexNAc and whether the final residue is GlcNAc or GalNAc.

In *Cj*, the enzyme following PglH is PglI. In *Cc*, there is a homolog assigned as PglI (38.2% sequence identity), and so our assumption was that the reaction catalyzed by *Cc* PglI follows that of PglH1. For *Cj* and *Cc* PglI, as well as PglI homologs, the canonical GT-A DXD motif is a conserved DDDD sequence. The AlphaFold-generated models of the *Cj* and *Cc* enzymes predict a classic GT-A fold consisting of an N-terminal Rossmann-like domain and a variable C-terminal domain (**Figure S1**) (33,34). The C-terminal domain of PglI as modeled by AlphaFold3, is an all α-helical domain that interacts with the membrane, as predicted using the PPM server (33,35), allowing access to the membrane-resident undecaprenyl diphosphate-sugar acceptor.

Both *Cc* and *Cj* PglI were heterologously expressed and purified in detergent to modest purity (**Figure S1**). Initial activity of the PglIs from *Cj* and *Cc* was assessed via radioactivity-based biochemical assays. In these assays, the transfer of a tritiated sugar from a UDP-[^3^H]sugar donor to the polyprenyl-diphosphate-linked acceptor is monitored. The undecaprenyl diphosphate-[^3^H]-sugar product is extracted through partitioning of a chloroform/methanol/water mixture and measured in percent conversion in dpm, which represents the organic layer radioactivity (undecaprenyl-linked) as a percentage of the total dpm for the reaction (organic and aqueous layer, wherein the aqueous layer contains unreacted UDP-[^3^H]sugars). The donor sugars evaluated included: [^3^H]Glc, [^3^H]GlcNAc, [^3^H]GalNAc, and [^3^H]Gal. The appropriate acceptors (PglH product for *Cj* and PglH1 product for *Cc*) were biochemically synthesized using the preceding enzymes in the pathway.

Consistent with the literature, we observed that *Cj* PglI preferentially installs a [^3^H]Glc in the PglH product (**Figure 2**, GalNAc_5_-diNAcBac-PP-Und). There is negligible activity (35.2 to 377.6 DPM, 0.1 to 0.8% fraction organic relative to total counts) with all other sugars compared to Glc addition (4646.5 DPM, 15.5% fraction organic). In contrast, *Cc* PglI exhibits significant activity for the transfer of either Glc(3267.6 DPM, 11% fraction organic) or GlcNAc (4571.2 DPM, 8.6% fraction organic) to the PglH1 product (**Figure 2**, GlcNAc_2_-GalNAcA-GalNAc-diNAcBac-PP-Und). It was unexpected that the PglI would append a GlcNAc, as there are significant structural differences between GlcNAc and Glc and GlcNAc has not previously been predicted to be a substrate for PglI, based on the *Cj* enzyme role of inserting a β-1,3-linked Glc. It appears to be a unique feature of *Cc* PglI, as *Cj* PglI did not exhibit the same promiscuity. It is also worth noting that *Cc* PglI does display some promiscuity with regard to the acceptor provided— *Cc* PglI is able to transfer Glc to the *Cj* PglH acceptor, GalNAc_5_-diNAcBac-PP-Und, despite the difference in sugar-chain lengths (**Figure 2**). It is possible that the promiscuity of *Cc* PglI increases diversity of the final glycoconjugates, enabling structural variation that promotes glycan evolution and hence, bacteria survival. The ability of *Cc* PglI to insert a GlcNAc matches the analysis of the final *Cc* glycoconjugate (20), though it is unexpected that this GlcNAc addition would be performed by PglI instead of PglH1 or PglH2 appending additional GlcNAc monosaccharides, as PglH performs a processive addition of GalNAc in *Cj*.

**Figure 2.**
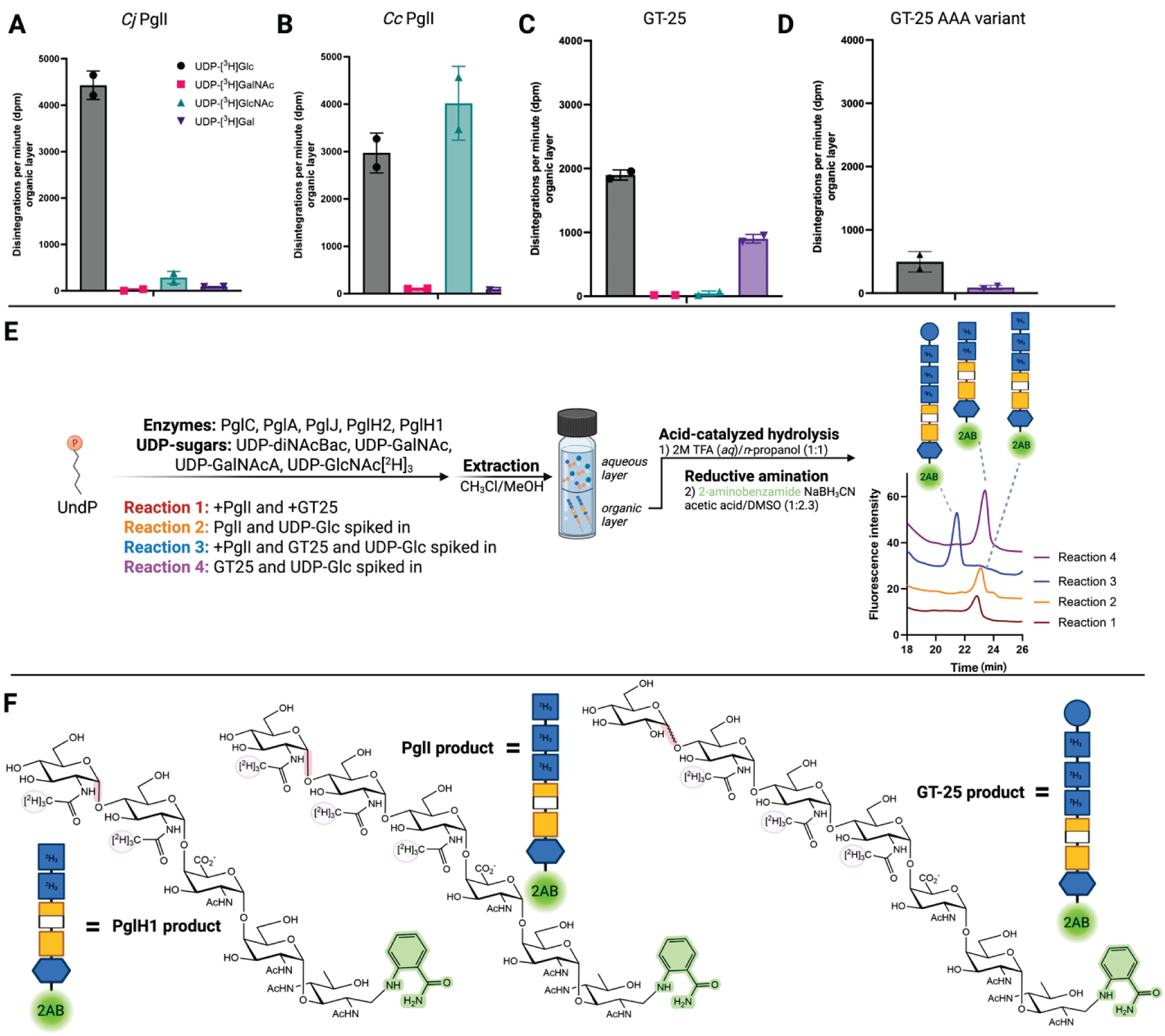
Sugar specificity of PglI and GT-25. Radioactivity analysis of sugar preferences of: (**A**) CjPglI with (GalNAc)_5_-diNAcBac-UndPP; (**B**) CcPglI with GlcNAc-GlcNAc-GalNAc-diNAcBac-UndPP; (**C**) GT-25 with GlcNAc-GlcNAc-GlcNAc-GalNAc-diNAcBac-UndPP; and (**D**) GT-25 AAA variant with GlcNAc-GlcNAc-GlcNAc-GalNAc-diNAcBac-UndPP. (**E**) Liquid chromatography traces of different reactions with proposed glycans. Reaction components are listed in **Table 1**. Reaction 1 includes PglC through GT-25 without UDP-Glc; reaction 2 includes PglC to PglI without UDP-Glc; reaction 3 includes PglC to GT-25 with UDP-Glc; and reaction 4 includes PglC to GT-25 and omits PglI. (**F**) Chemical drawings of the three products observed in **E**.

The predicted *Cc* glycoconjugate includes an additional Hex residue at a branching position, reminiscent of the position of the carbohydrate addition performed by *Cj* PglI. As *Cc* PglI appears to be attaching a GlcNAc, that Hex is potentially appended by a different enzyme. It is possible that PglI performs two glycosyl transfers, but this was not observed in our experimental analysis. When we analyzed the genome neighborhood network for several different species of *Camplyobacter*, we identified a gene, *gt-25*, preceding *pglI* in the operon, annotated as encoding a glycosyltransferase family-25 enzyme (**Figure 1A**). This gene was typically found downstream of *pglk* in the operon context, suggesting that it plays a role in the *pgl* pathway. Glycosyltransferase family-25 enzymes belong to the GT-A superfamily and include approximately 14,000 members (36). These enzymes are found primarily in bacterial lipopolysaccharide biosynthesis and transfer sugars onto the growing glycan chain (37). Enzymes in this class contain the hallmark DXD motif of the GT-A family and are predicted to catalyze carbohydrate transfer via an inverting mechanism. In GT-25, the DXD motif is slightly modified to E^90^D^91^D^92^ but aligned in primary and tertiary structure with the corresponding motif in PglI. The structure of GT-25 predicted by AlphaFold shows the GT-A family fold, composed of a Rossmannoid N-terminal domain, followed by a variable C-terminal domain and shows some similarity to the overall structure of PglI (**Figure S2**).

GT-25 and the GT-25 AAA variant were heterologously expressed and purified in detergent to moderate purity (∼80%, **Figure S2**). NanoDSF was employed to assess sugar donor preference of wild-type GT-25 through thermal stabilization (**Figure S3**). There was moderate stabilization in the presence of all the UDP-sugar donors as opposed to the sample without added ligand, but the greatest stabilization occurred in the presence of either UDP-Glc or UDP-Gal. To determine if GT-25 follows PglI sequentially in the biosynthetic pathway, we utilized a radioactivity-based assay (**Figure 2**). The acceptor (GlcNAc-GlcNAc-GlcNAc-GalNAcA-GalNAc-diNAcBac-PP-Und) was biochemically synthesized using the biosynthetic enzymes PglC through PglI. GT-25 was able to transfer UDP-Glc or UDP-Gal to the PglI product (**Figure 2**), however, there was greater activity with UDP-Glc (1899.4 DPM, 3.5%), than UDP-Gal (901.2 DPM, 1.6%). We also evaluated the activity of the GT-25 AAA variant and found minimal activity (1.25% in the organic fraction for Glc, 0.10% in the organic fraction for Gal), confirming that the EDD motif plays a role in binding and/or catalysis of the glycosyltransferase reaction (**Figure 2**).

### Evaluating the interactions of PglI and GT-25

It is likely that the enzymes involved in the *pgl* pathway colocalize and form a metabolon, to more efficiently facilitate glycoconjugate biosynthesis and ensure substrate flux. This is particularly important because UndP concentrations are very low (<0.1 mol%) in bacterial membranes (38,39). Our laboratory has obtained evidence that the early enzymes in the pathway (*Cj* PglC, PglA, and PglJ) colocalize (39). In addition, during the chemoenzymatic syntheses, we observed an increase in the amount of overall product when a greater number of sequentially acting enzymes were included, suggesting product transfer between PGTs/GTs. To probe this, PglI activity was assessed with UDP-GlcNAc donor and the pentasaccharide acceptor and UDP production monitored in the presence and absence of GT-25 and the GT-25 AAA variant (**Figure S5**). GT-25 has negligible activity with UDP-GlcNAc and this acceptor, so the effect of mass action can be ruled out. In the presence of GT-25 or the GT-25 AAA variant, increased PglI activity was observed, producing 1.6 µM and 0.4 µM more UDP, respectively, over the time course of the reactions (**Figure S5**). The enhancement in UDP production by PglI in the presence of GT-25 provides support for substrate channeling and interaction of the two proteins *in vivo*. It is noted that in a pull-down assay using detergent-solubilized enzymes, PglI did not capture either PglH1 or GT-25 under the experimental conditions (**Figure S4**). However, it is anticipated that the native co-localization in membranes would limit the degrees of freedom, lowering the entropic penalty and enhancing affinity.

### Confirmation of glycan composition with 2-AB labeling and mass spectrometry

The glycan composition of the undecaprenyl diphosphate-linked products was confirmed by isolating the products by extraction, followed by hydrolysis and 2-aminobenzamide (2-AB) labeling, allowing for preliminary characterization of glycans by HPLC and low-resolution mass spectrometry (14,23,27). After the enzymatic reaction, undecaprenyl diphosphate-glycans were extracted from the product mixture with chloroform and the water-soluble, unreacted UDP-sugar substrates were removed with aqueous washes. The organic layers were concentrated, then hydrolyzed with 2 M trifluoroacetic acid (TFA), to remove the glycans from the polyprenyl diphosphate. Following this, the reducing sugar end was labeled with 2-AB through reductive amination with NaBH_3_CN. The mixture was purified by HPLC using fluorescence detection. Peaks were collected and a preliminary analysis performed by negative-mode electrospray ionization mass spectrometry (ESI-MS) using a single-quadrupole mass analyzer. For the larger glycans, the doubly-charged species [M – 2H]^2-^ had higher abundance than the singly-charged species [M - H]^1-^ (**Table S1**). Distinction between GalNAc or GlcNAc additions was facilitated by utilization of [^2^H_3_] GlcNAc.

As discussed below, assignments were later verified by high resolution positive-ion nanoESI-MS and HCD MS/MS of [M + X]^1+^ and M + 2X]^2+^ (X = H, Na and/or K) performed on a tribrid Orbitrap instrument that has mass accuracy well below 10 ppm (**Table S2**). As demonstrated in **Figures 3, 4** and **Figures S6 - S9**, the linkage positions and the presence of isomeric products were established by EED tandem mass spectrometry of these species, performed on a hybrid Orbitrap instrument modified by addition of an Omnitrap [18].

**Figure 3.**
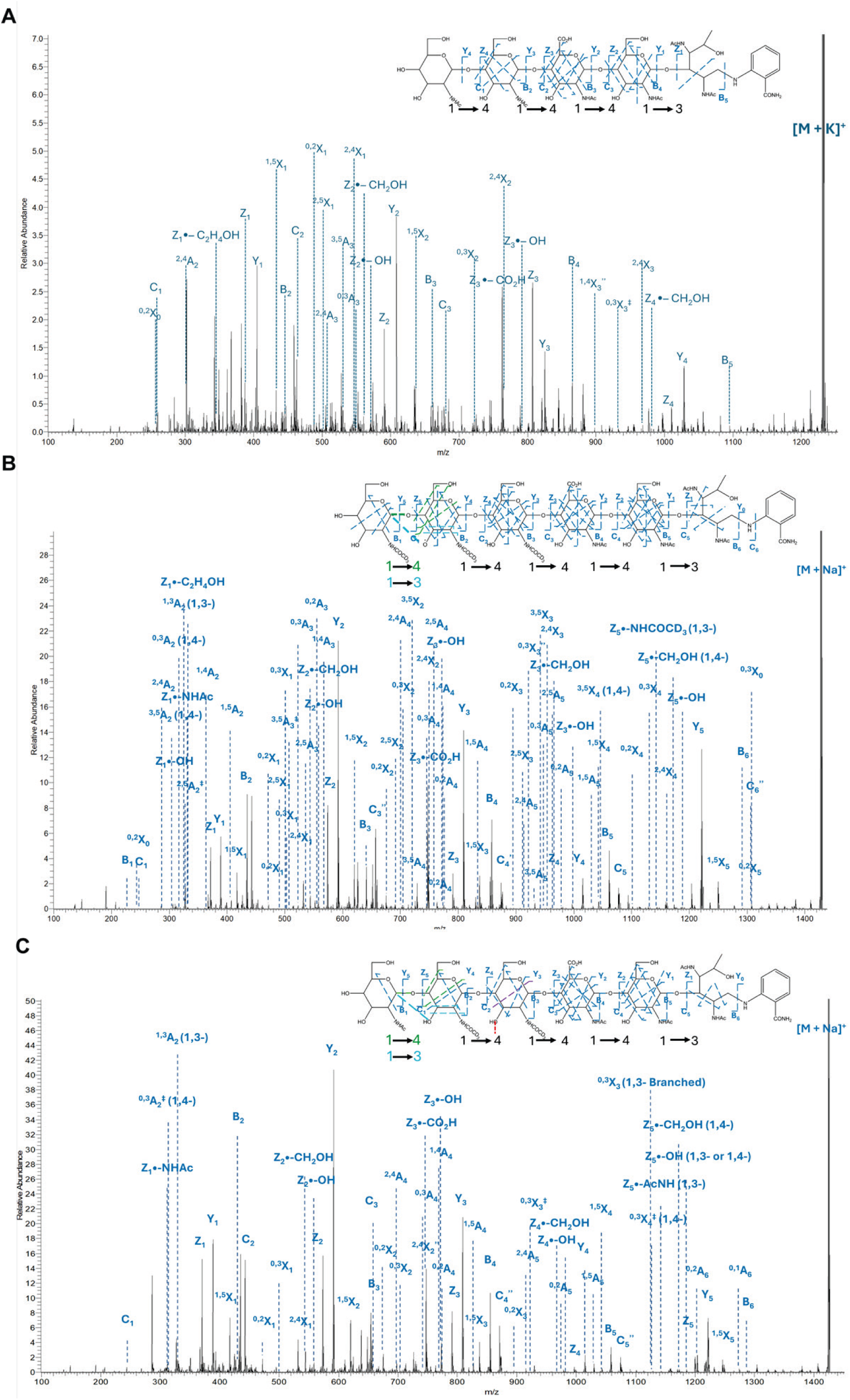
EED mass spectrometry analysis of PglH1 and PglI products. (**A**) EED MS/MS spectrum of the PglH1 product [M + K]^1+^.*m/z* 1237.4849 (**B**) EED MS/MS spectrum of the [^2^H]_3_ PglI product [M + Na]^1+^ *m/z* 1427.6157.(**C**) EED MS/MS spectrum of the [^2^H]_2_[H]_3_ PglI product [M + Na]^1+^ *m/z* 1424.5934.

In agreement with previous work, the observed PglH1 product was [^2^H_3_]GlcNAc-[^2^H_3_]GlcNAc-GalNAcA-GalNAc-diNAcBac-2-AB (**Figure 2E**) (23). The same reaction was carried out in the presence of PglI and GT-25, with all sugars present save for UDP-Glc (**Table S1**). As shown in **Figure S6**, a peak was present at *m/z* 1403.4 [M - H]^1-^ in the negative-ion mass spectrum. This most likely corresponds to the addition of a GlcNAc (as PglI does not utilize GalNAc, GalNAcA, or diNAcBac). The product of this reaction would be [^2^H_3_]GlcNAc-[^2^H_3_]GlcNAc-[^2^H_3_]GlcNAc-GalNAcA-GalNAc-diNAcBac-2-AB (**Figure 2E**). This is consistent with the experiment using radiolabeling, which suggested that PglI was able to append GlcNAc as well as Glc. Because the radiolabeled experiments demonstrated that GT-25 is unable to append GlcNAc, PglI is the likely source of the GlcNAc addition. The experiment was repeated excluding GT-25 and including UDP-Glc and spiking in PglI after 60 min; the major species observed was [^2^H_3_]GlcNAc-[^2^H_3_]GlcNAc-[^2^H_3_]GlcNAc-GalNAcA-GalNAc-diNAcBac-2-AB (**Figure 2E**). To confirm that GT-25 does not act on the PglH1 product, the sequential enzymes PglC through PglH1 were combined and allowed to react before spiking in GT-25 and UDP-Glc; the only abundant peak observed in the mass spectrum corresponded to the PglH1 product (**Table S1, Figure 2E**). These experiments confirmed the order of the PglI and GT-25 glycan additions as well as identified PglI as the source of the final GlcNAc residue on the growing glycan chain.

In the PglH1 structure, two residues were identified as playing a crucial role in stereospecific-donor substrate fidelity (S258 and N259) (23). The S258P/N259T variant of PglH1 appends a GalNAc instead of GlcNAc to the PglH2 product. We tested the PglI acceptor specificity by assaying the ability of PglI to append a GlcNAc to the PglH1 S258P/N259T product, GalNAc-[^2^H_3_]GlcNAc-GalNAcA-GalNAc-diNAcBac-PP-Und. A singly-charged peak was observed at *m/z* 1400.6, corresponding to the [M - H]^-^ calculated for [^2^H_3_]GlcNAc-GalNAc-[^2^H_3_]GlcNAc-GalNAcA-GalNAc-diNAcBac-2-AB (**Figure S7**). This was not unexpected, as *Cc* PglI was able to append Glc to the *Cj* hexasaccharide acceptor with a non-reducing end GalNAc, rather than a reducing-end GlcNAc. Notably, the *Cj* acceptor for PglI is one glycan unit longer than the *Cc* pentasaccharide acceptor. Overall, this indicates that PglI is promiscuous both in the donor sugar identity and non-reducing end acceptor identity as well as the length of the acceptor.

Finally, PglC through PglH1 were sequentially reacted and then PglI, GT-25, and UDP-Glc were added simultaneously. The most abundant peak in the mass spectrum was a doubly-charged signal at *m/*z 782.5. This would correspond to the [M - 2H]^2-^ of Glc-[^2^H_3_]GlcNAc-[^2^H_3_]GlcNAc-[^2^H_3_]GlcNAc-GalNAcA-GalNAc-diNAcBac-2-AB. This finding is consistent with the chemoenzymatic syntheses and radiolabeled experiments where it was posited that PglI appends a GlcNAc and GT-25 appends a Glc. However, the low resolution MS1 spectra provide only nominal molecular weight information and did not define the locations of the sugars newly added by PglI and GT-25. PglI in *Cj* appends a branching β-1,3-linked Glc to the penultimate glycan at the non-reducing end. The GT-25 family is annotated to transfer glycan residues with inversion of stereochemistry. The low resolution MS1 spectra supported the model where PglI appends a GlcNAc to the PglH1 product and GT-25 acts on the product of PglI and likely appends a Glc. The advanced tandem MS methods discussed below were required to fully define the structural details of the products of each enzymatic reaction.

### Identification of glycosidic linkage positions using EED tandem mass spectrometry

High resolution MS1 spectra of the AB derivatives of the glycan products provided the accurate mass measurements that are presented in **Table S2** to verify the compositions. Because MS1 spectra provide molecular weight information but no structural details, fragmentation of the molecular species and further stages of MS analysis (tandem MS, MS/MS) are needed to gain information on the glycan sequence and linkage positions. Collisional activation (CAD or CID, HCD) of glycans generates primarily cleavages of the glycosidic linkages, and thus can help define the glycan topology, but yields little information on the linkage positions. Electron-based fragmentation methods (ExD) generate extensive cleavages within the glycan rings that provide rich information on both the glycan topology and the linkage positions. We have found that EED is the most useful ExD option for glycans and glycoconjugates (40), and have determined that the sensitivity and speed of the hybrid Q Exactive HF-Omnitrap system significantly exceed the results that can be obtained with EED on an FT-ICR system (31), and therefore chose the Q Exactive HF-Omnitrap system for detailed study of the *Campylobacter N*-linked glycans.

First, the products generated by PglJ and PglH2 were characterized by EED MS/MS. The EED spectrum was obtained for the [M + Na]^1+^ *m/z* 809.3156 of the 2-AB–labeled PglJ product (GalNAcA–GalNAc–diNAcBac-2AB) (**Figure S8)**. Fragmentation yielded extensive glycosidic bond cleavage ions (B, C, Y, and Z series) alongside cross-ring fragments from the A- and X-ion series. The glycosidic fragments established the sequence GalNAcA–GalNAc-diNAcBac-2AB. Importantly, the detection of cross-ring ions, including ^3,5^X_1_, ^3,5^A_2_, and ^2,4^X_1_, excluded the presence of 1→4 linkages (32), whereas the observation of the ^0,2^X_n_ ion series supported a 1→3 linkage. Collectively, these data defined the PglJ glycan product as GalNAcA1→4GalNAc1→3diNAcBac.

The EED spectrum was obtained for the [M + Na]^1+^ *m/z* 1012.3999 of the 2-AB-labeled PglH2 product derived from the PglJ intermediate (**Figure S9)**. The presence of diagnostic cross-ring fragments, including ^3,5^X_2_, ^3,5^A_2_, ^2,4^X_2_, and ^2,4^A_2_ ions, indicates that the GlcNAc residue is attached via a 1→4 linkage. Therefore, the detailed structure of the PglH2 glycan product was determined to be GlcNAc1→4GalNAcA1→4GalNAc1→3diNAcBac.

The EED MS/MS spectrum of the [M + K]^1+^ *m/z* 1237.4849 of the 2-AB derivative of the PglH1 product was obtained (**Figure 3A)**. The majority of fragment ions correspond to reducing-end fragments, including complete B, C, Y, Z, and ^1,5^X-ion series. In addition, several linkage-dependent cross-ring fragments are observed. The presence of ^3,5^X, ^3,5^A, ^2,4^X and ^2,4^A ions in the spectrum rules out 1→4 linkages (32), whereas the presence of ^0,2^X and ^0,3^X ion series in the spectrum can be used to define its 1→3 linkage. Meanwhile, several secondary fragment ions generated by EED, such as Z^•^-CH_2_OH and Z^•^-OH further facilitate the determination of glycan linkages. The paired observation of Z^•^-CH_2_OH and Z^•^-OH ions can be used to support 1→4 linkages, whereas the presence of Z^•^-C_2_H_4_OH ion is indicative of a 1→3 linkage. Taken together, the detailed structure of the PglH1 glycan product was determined to be GlcNAc1→4GlcNAc1→4GalNAcA1→4GalNAc1→3diNAcBac.

Based on the PglH1 product, PglI was added to the sample together with UDP-GlcNAc[^2^H]_3_. The presence of ^3,5^A_2_, ^0,3^A_2_, and ^3,5^X_4_ ions in its spectrum (**Figure 3B**) indicates that the major component has GlcNAc attached to the PglH1 product though a 1→4 linkage catalyzed by PglI. Moreover, the observation of ^1,3^A_2_ and ^1,3^X_4_ ions indicates that a minor component is present in which GlcNAc is linked to the PglH1 product via a 1→3 linkage, also catalyzed by PglI. To differentiate between the linear glycan topology and the branched structures produced PglI, UDP-GlcNAc and UDP-GlcNAc were added separately to the sample. The EED MS/MS spectrum was obtained for the [M + Na]⁺ *m/z* 1424.5934 of the PglI products resulting from treatment with UDP-GlcNAc (**Figure 3C)**. The observed ^0,3^A_2_, ^0,3^X_4_ and Z ^•^-CH_2_OH fragments indicate that the major component corresponds to GlcNAc attachment to the PglH1 product via a 1→4 linkage catalyzed by PglI. Moreover, the observation of the ^1,3^A_2_ and Z ^•^-AcNH fragments shows that a minor component is present in which GlcNAc is linked to the PglH1 product via a 1→3 linkage appended by PglI. Furthermore, the presence of the ^0,3^X_3_ fragment ion indicates a small proportion of the products contain GlcNAc attached though a 1→3 linkage to the penultimate glycan residue at the non-reducing end of the PglH1 product. Overall, the PglH1 product was determined to have a structure containing three 1→4 linkages and one 1→3 linkage (read from the non-reducing end to the reducing end). The PglI product is predominantly linear, with a small proportion of branched structures. The terminal GlcNAc residues at the non-reducing end of the PglI product have both 1→3 and 1→4 linkages.

Additionally, PglI was added to the PglH1 product together with UDP-Glc. The resulting products were subjected to EED MS/MS analysis, and the MS/MS spectrum obtained for the [M + Na]^1^⁺ *m/z* 1405.5519 is shown in **Figure S10**. Similarly, Glc is predominantly linked to the PglH1 product via a 1→4 linkage catalyzed by PglI.

GT-25, the GT downstream in the pathway, was added to the sample together with UDP-Glc, with the result that both the linear structure and branched structures were determined in the GT-25 products. For each of the isomeric structures the EED spectrum of [M + Na + H]^2+^ *m/z* 790.8072 was recorded (**Figure 4**). In the EED spectrum of the linear structure (**Figure 4A**) the presence of ^3,5^A_2_ and ^1,4^A_2_ ions indicates that Glc is predominantly attached to the PglI product though a 1→4 linkage catalyzed by GT-25. Accordingly, the structure assigned to the most abundant GT-25 glycan product is: GlcNAc1→4Glc1→4 GlcNAc1→4GlcNAc1→4GalNAcA1→4GalNAc1→3diNAcBac. The EED MS/MS spectra of the branched GT-25 products (also having [M + Na + H]^2^⁺ *m/z* 790.8072) were also obtained (**Figures 4B** and **4C)**. In the spectrum shown in **Figure 4B**, the presence of ^3,5^A_2_, ^0,3^A_2_ and ^1,3^A_2_ ions indicates that the Glc is attached to the PglI product though a branching 1→3 linkage catalyzed by GT-25, yielding the glycan structure Glc1→4(GlcNAc1→3)GlcNAc1→4GlcNAc1→4GalNAcA1→4GalNAc1→3diNAcBac. In contrast, in **Figure 4C**, the observation of ^3,5^A_2_, ^0,3^A_2_ and ^1,3^A_2_ ions indicate that the Glc is attached to the PglI product though a branching 1→3 linkage, that generates the glycan Glc1→4(GlcNAc1→3)GlcNAc1→4GlcNAc1→4GalNAcA1→4GalNAc1→3diNAcBac.

**Figure 4.**
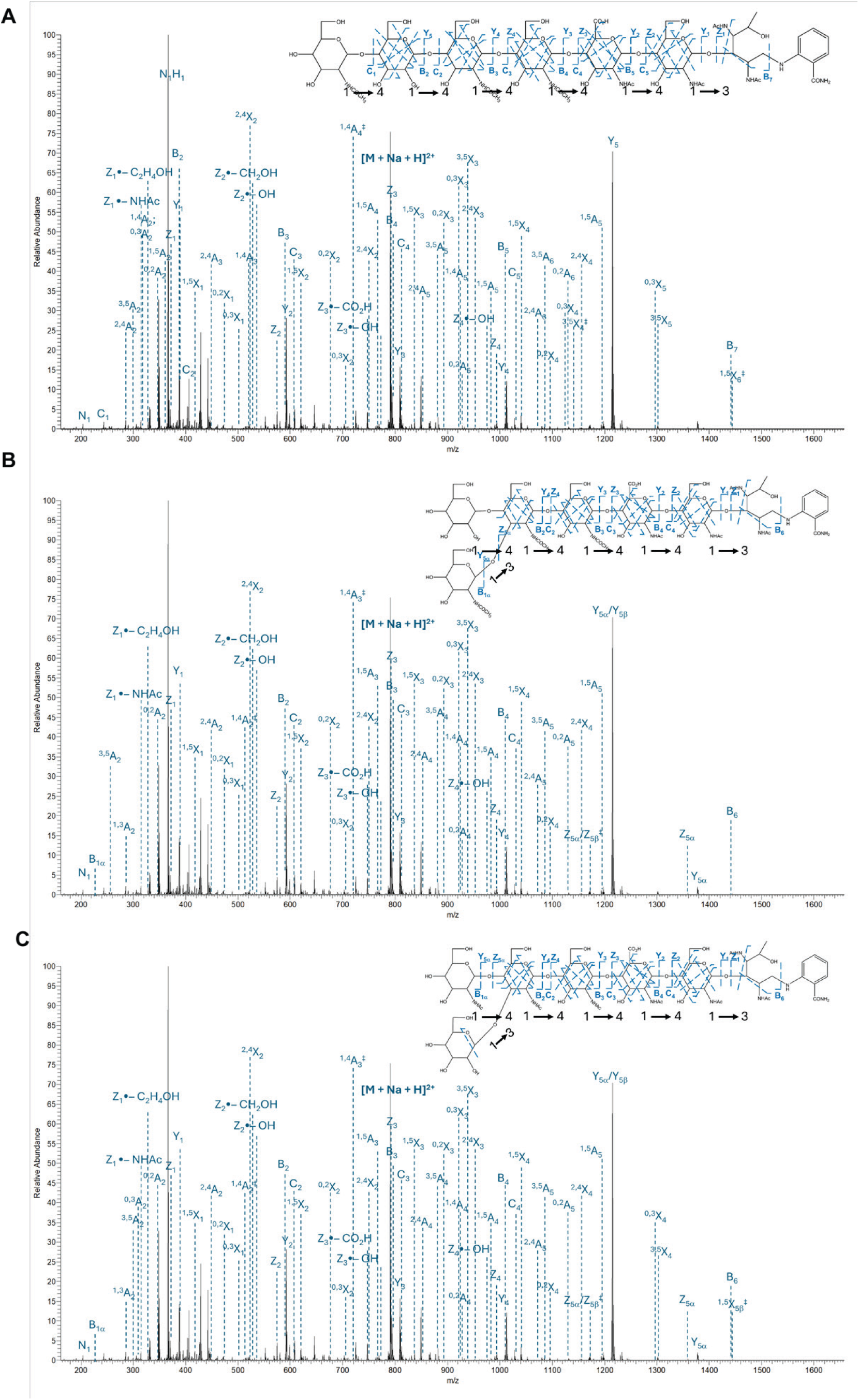
EED mass spectrometry analysis of GT25 products. (**A**) EED MS/MS spectrum of [M + Na + H]^2+^ *m/z* 790.8072 of the GT-25 product assigned to the structure GlcNAc1→4Glc1→4GlcNAc1→4GlcNAc1→4GalNAcA1→4GalNAc1→3-diNAcBac-2-AB. (**B**) EED MS/MS spectrum of [M + Na + H]^2+^ *m/z* 790.8072 of the GT-25 product assigned to the structure Glc1→4(GlcNAc1→3)GlcNAc1→4GlcNAc1→4GalNAcA1→4GalNAc1→3-diNAcBac-2-AB. (**C**) EED MS/MS spectrum of [M + Na + H]^2+^ *m/z* 790.8072 of the GT-25 product assigned to the structure GlcNAc1→4(Glc1→3)GlcNAc1→4GlcNAc1→4GalNAcA1→4GalNAc1→3-diNAcBac-2-AB.

Collectively, these results demonstrate that GT-25 is capable of catalyzing both linear 1→4 and branching 1→3 glucosyl transfer reactions on the PglI product.

## Conclusions

In this work, we report the biochemical characterization of the two final glycosyltransferases in the *Cc pgl* pathway and identify the resulting glycoconjugates with the glycan residue, position, and linkage position assignments for each of the carbohydrate addition steps. The substrate preferences for PglI and GT-25 were determined through nanoDSF analysis, kinetic studies, radiolabeled sugar donor assays, and mass spectrometry. Detailed definition of the 2-AB glycan derivatives was achieved by using an advanced mass spectrometry approach that employed the newly developed EED fragmentation method to enable assignment of the linkage positions in addition to the glycan sequences, even in cases where heterogeneous products were formed. The detailed structures of the glycans produced from PglJ, PglH2, PglH1 were fully determined. Notably, PglI was found to be able to transfer either of two sugars under physiologically relevant concentrations. PglI in most Campylobacter species is annotated to transfer Glc (14,19). Our findings show that in *Cc*, PglI is capable of catalyzing transfer of either GlcNAc or Glc, but when both sugars are available, PglI predominantly transfers GlcNAc. Notably, PglI is able to transfer GlcNAc or Glc to the linear chain through both a 1→4 and 1→3 linkage, though 1→4 is the preferred product. In terms of anomeric configuration, current assignments are based on protein fold and sequence identity, where the *Cc* glycosyltranferases are predicted to make the same linkages as the orthologs in *Cj*. PglI is an inverting glycosyltransferase that installs a β-linked glucose residue and likewise, GT-25 family glycosyltransferases are generally associated with inverting reactions that produce β-configured glycosidic linkages (41). Although currently the anomeric stereochemistry needs to be defined independently, there is promise that advances in mass spectrometry will ultimately enable direct assignment.

GT-25 was formerly overlooked as a contributor to glycan assembly in *Cc*. Previously, it was suggested that GT-25 may play a role in sugar modification to produce a 234 Da sugar, proposed to be the monosaccharide glucolactilic acid (19). However, another study (20) did not find the 234 Da sugar in the final glycan product. This modified sugar has been consistently found in *Campylobacter* that contain *pgl* operons with a pyruvyl transferase gene. The pyruvyl transferase in *E. coli,* MurA, is known to modify UDP-*N*-acetyl-α-D-glucosamine directly before further sugar elaboration for peptidoglycan assembly (42). However, the absence of an orthologous gene to MurA suggests that this highly-modified sugar is not generated in this strain of *Cc.* We have demonstrated that GT-25 acts on the PglI product and is the source of the appended Glc, although it can transfer both Glc and Gal.

GT-25 is of interest as it is able to append Glc to the product of PglI in three different ways (in order of abundance): linear 1→4; branching 1→3; and linear 1→4 when PglI has appended a linear 1→3. The specificity of PglI and GT-25 reported here, together with that of H2 and H1 for Glc and/or GlcNAc (23), defines the structure of the glycan of *Cc*, which is in a subgroup of Group II organisms (including *Campylobacter curvus* and *Campylobacter rectus*) (4). Along with those characterized in Group II, our findings show that GlcNAc predominates at the non-reducing end of the ultimate glycan. This usage contrasts with that of the Group I species (such as *Cj* and *Campylobacter coli*), where GalNAc predominates (4,19). UDP-GalNAc is generated through reversible epimerization of UDP-GlcNAc, therefore, its intracellular abundance is lower and is inherently dependent on the size and flux of the UDP-GlcNAc pool (43). It is possible, therefore, that *N*-glycan composition may, in part, be steered by the availability of sugar donor substrates.

The observation that GT-25 appends Glc at different branching points is significant, especially when considered in the context of the promiscuity of PglI, which transfers GlcNAc or Glc to different acceptor sites. It appears that PglI and GT-25 allow the diversification of the final glycoconjugate in the Group II *Cc*. Overall, *Campylobacter* Group I and Group II species possess strikingly different patterns of glycan diversity and ecological adaptation. In Group I species, including *Cj* and *Camplyobacter lari* there is conservation of the core hexasaccharide (4,19). The diversity observed in these studies of *Cc* with respect to acceptor-substrate recognition and the actions of PglI and GT-25 is consistent with the extensive variation in glycan length, composition, and chemistry displayed in Group II species. The nonreducing-end sugars are key determinants of adaptive and innate immune recognition (44,45), thus host recognition mechanisms likely impose selective constraints that shape the evolution of *N*-glycan diversity (18). This model is supported by the chemically distinct C6-oxidized sugar, GalNAcA, observed in Group II species, including *Cc* 13826 and *Cc* 33237 (24), which occupy the specialized host niche of the human oral cavity. The ability of the *pgl* pathway to adapt and produce a variety of glycoconjugates could impact bacterial survival and pathogenicity (6) and underscores a role for immune-mediated selection in the evolution of *N*-linked oligosaccharide biosynthesis in *Campylobacter*.

An additional facet to the observed diversity is that there may be a mechanism allowing the bacteria to differentiate between the multiple glycoforms such that only selected forms will be ultimately *N*-linked. When analyzing the *pgl* operons, there are several instances of two PglB *N*-oligosaccharyltransferase enzymes (**Figure 1A**) (20). Many of the operons with genes encoding two PglB orthologs also encode GT-25, so potentially, the PglB may play a role in distinguishing which of the final glycoforms are displayed. Further *in vivo* study is needed to understand why multiple glycoforms may be observed and the role that the two PglB enzymes play in these biosynthetic pathways.

The definition of the role of the individual *pgl* enzymes in *Cc* and the identity of the final *N*-linked glycoconjugate revealed unanticipated differences in carbohydrate composition. Specifically, we note that a far higher proportion of GlcNAc is observed in the Group II glycan at the non-reducing end, which would have a dramatic effect on interactions with cognate receptors (45,46). This study expands the understanding of the Group II glycans and the selection of their glycoforms which could be mediated by both metabolic and evolutionary pressures

## Supporting information

Supplement containing Tables S1 & S2 and Figs S1-S10

## Acknowledgments

We thank Dr. Soumi Ghosh for providing the pulldown assay protocol and Dr. Nemanja Vuksanovic for providing the *Cc* PglH1 S258P/N259T variant.

## TOC graphic

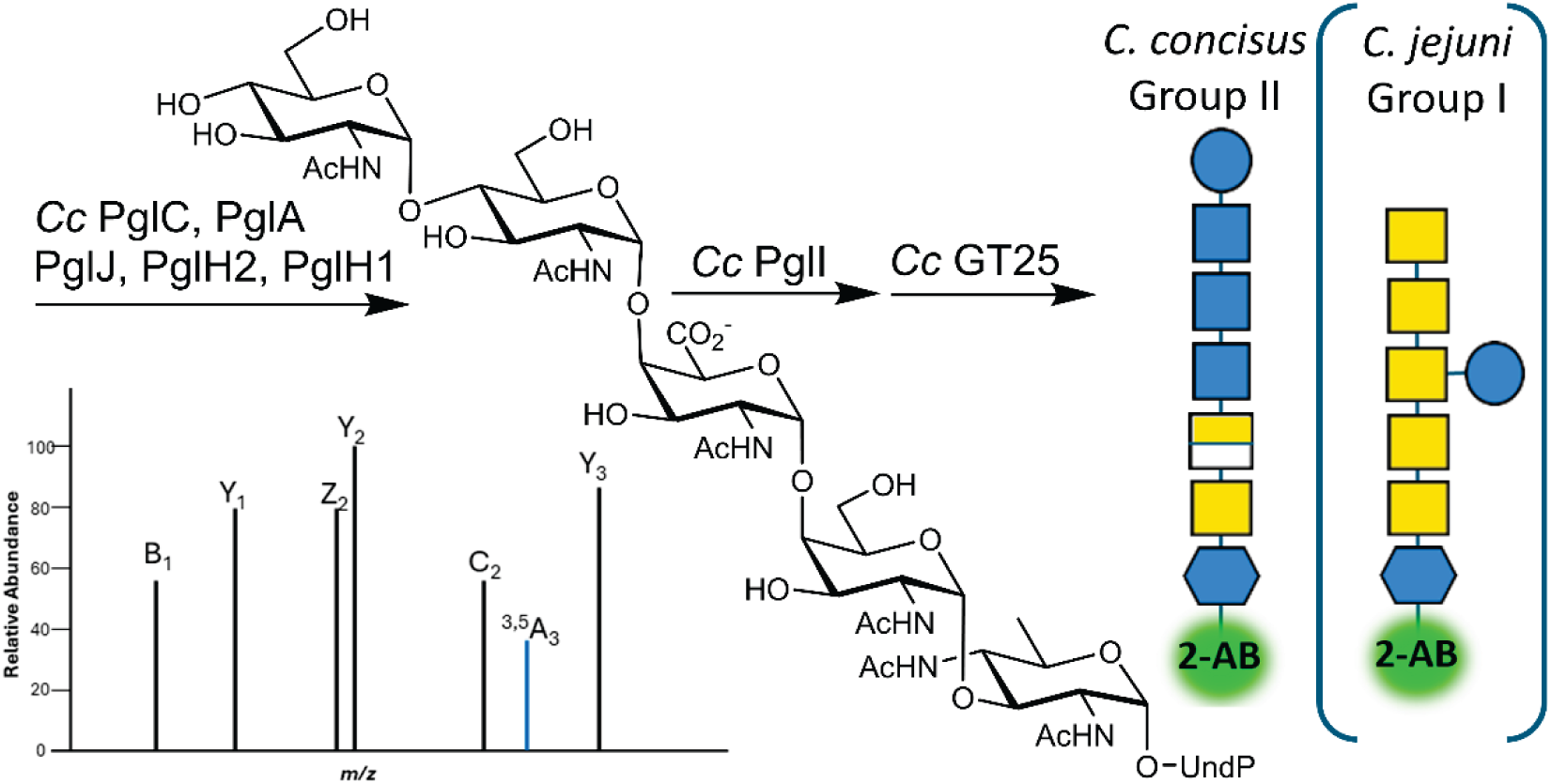

