## Supplement containing Tables S1 & S2 and Figs S1-S10 for "Glycoconjugate diversification in *Campylobacter concisus* is determined by two glycosyltransferases"

### Diversification of the glycoconjugate in *Campylobacter concisus* is governed by two glycosyltransferases

**Table S1.** Reactions run with experimental parameters and results from low resolution negative-ion ESI mass spectrometry measurements in the range *m/z* 100 to 1500.

| Reaction | Enzymes | Sugars | Predominant mass (*m/z*) | Expected mass (*m/z*) | Glycan |
|---|---|---|---|---|---|
| **1** | PglC, PglA, PglJ, PglH2, PglH1, PglI, GT-25 | UDP-Bac, UDP-GalNAc, UDP-GalNAcA, UDP-GlcNAc-[$^2H_3$] | 1403.4 | 1404.4 | $GlcNAc_3$-GalNAcA-GalNAc-diNAcBac-2AB |
| **2** | PglC, PglA, PglJ, PglH2, PglH1, PglI spiked in | UDP-Bac, UDP-GalNAc, UDP-GalNAcA, UDP-GlcNAc-[$^2H_3$], UDP-Glc spiked in | 1403.4 | 1404.4 | $GlcNAc_3$-GalNAcA-GalNAc-diNAcBac-2AB |
| **3** | PglC, PglA, PglJ, PglH2, PglH1, PglI and GT-25 spiked in | UDP-Bac, UDP-GalNAc, UDP-GalNAcA, UDP-GlcNAc-[$^2H_3$], UDP-Glc spiked in | 782.5 ($2^-$) | 1566.6 | Glc-$GlcNAc_3$-GalNAcA-GalNAc-diNAcBac-2AB |
| **4** | PglC, PglA, PglJ, PglH2, PglH1, GT-25 spiked in | UDP-Bac, UDP-GalNAc, UDP-GalNAcA, UDP-GlcNAc-[$^2H_3$], UDP-Glc spiked in | 1197.4 | 1198.2 | $GlcNAc_2$-GalNAcA-GalNAc-diNAcBac-2AB |
| **5** | PglC, PglA, PglJ, PglH2, PglH1 S258P/N259T, PglI | UDP-Bac, UDP-GalNAc, UDP-GalNAcA, UDP-GlcNAc-[$^2H_3$], UDP-Glc spiked in | 1400.6 | 1401.4 | GlcNAc-GalNAc-GlcNAc-GalNAcA-GalNAc-diNAcBac-2AB |

**Table S2.** Accurate mass values determined for intact molecular species via positive-ion high resolution mass spectrometry measurements

| Enzymes | Observed ions | Theoretical *m/z* | Observed *m/z* | Δ, ppm | Glycan |
|---|---|---|---|---|---|
| Pgl- C, A, J | $[M + Na]^{1+}$ | 809.3176 | 809.3156 | -2.4 | GalNAcA-GalNAc-diNAcBac-2AB |
| Pgl- C, A, J, H2 | $[M + Na]^{1+}$ | 1012.3970 | 1012.3999 | +2.9 | GlcNAc-GalNAcA-GalNAc-diNAcBac-2AB |
| Pgl- C, A, J, H2, H1 | $[M + Na]^{1+}$ | 1221.5140 | 1221.5164 | +1.9 | $GlcNAc_2$-GalNAcA-GalNAc-diNAcBac-2AB |
| Pgl- C, A, J, H2, H1, I | $[M + Na]^{1+}$ | 1427.6122 | 1427.6159 | +2.5 | $GlcNAc_3$-GalNAcA-GalNAc-diNAcBac-2AB |
| Pgl- C, A, J, H2, H1, I, GT-25 | $[M + 2Na]^{2+}$ | 801.7989 | 801.7989 | 0.0 | Glc-$GlcNAc_3$-GalNAcA-GalNAc-diNAcBac-2AB |

**SI Figures**

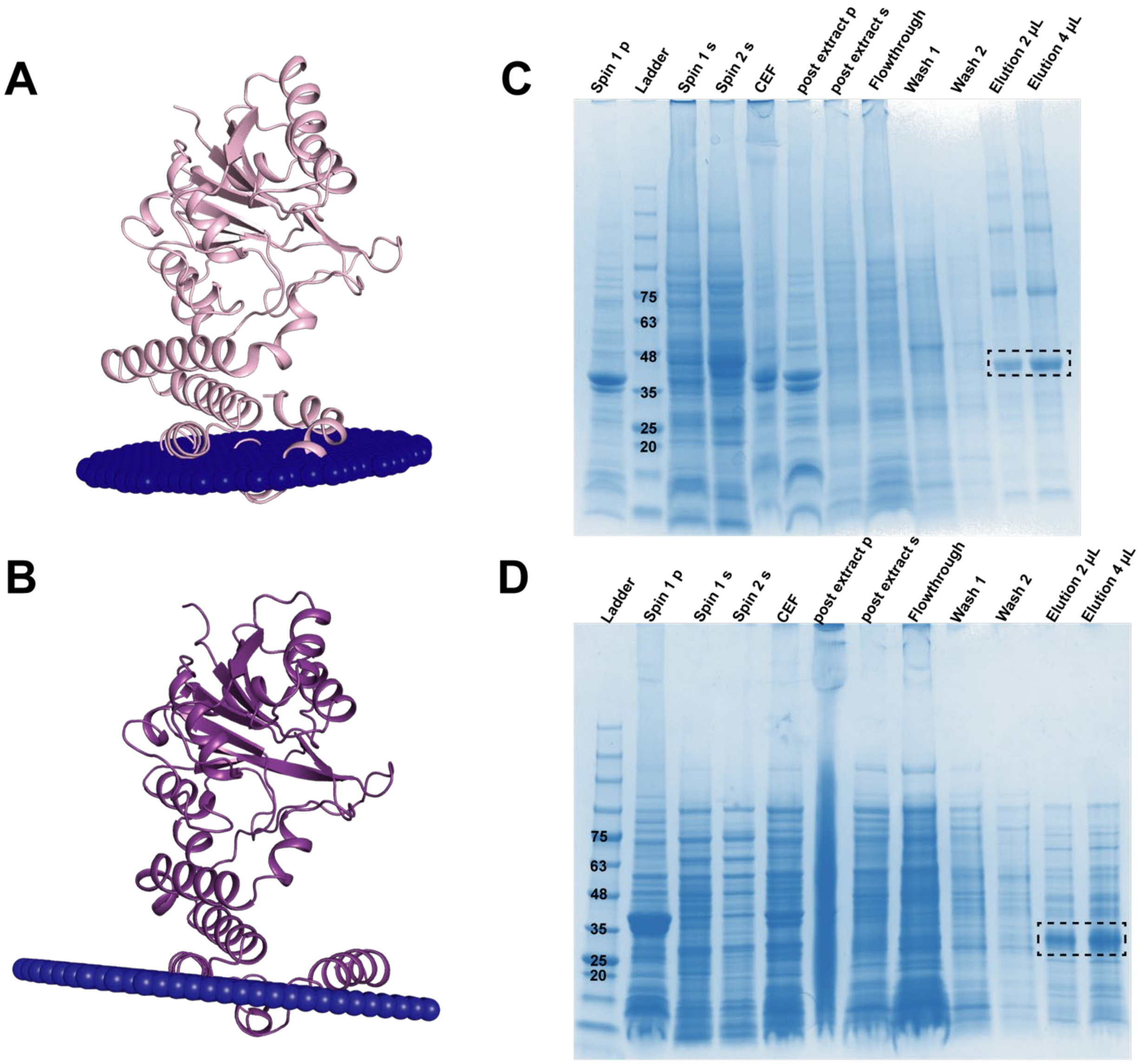


**Figure S1**. Comparison of *Cj* PglI and *Cc* PglI. Sequence alignment between the two proteins shows 38.2% identity. (A) AlphaFold model of *Cj* PglI docked into the membrane (shown as blue spheres) by the PPM server. (B) Alphafold model of *Cc* PglI docked into the membrane (shown as blue spheres) by the PPM server. The RMSD of the PglI ortholog AlphaFold structures is 1.4 Å. (C) SDS-PAGE of *Cj* PglI purification steps. Pellet (p) and supernatant (s) are abbreviated. Volume listed represents the amount loaded. (D) SDS-PAGE of *Cc* PglI purification steps.

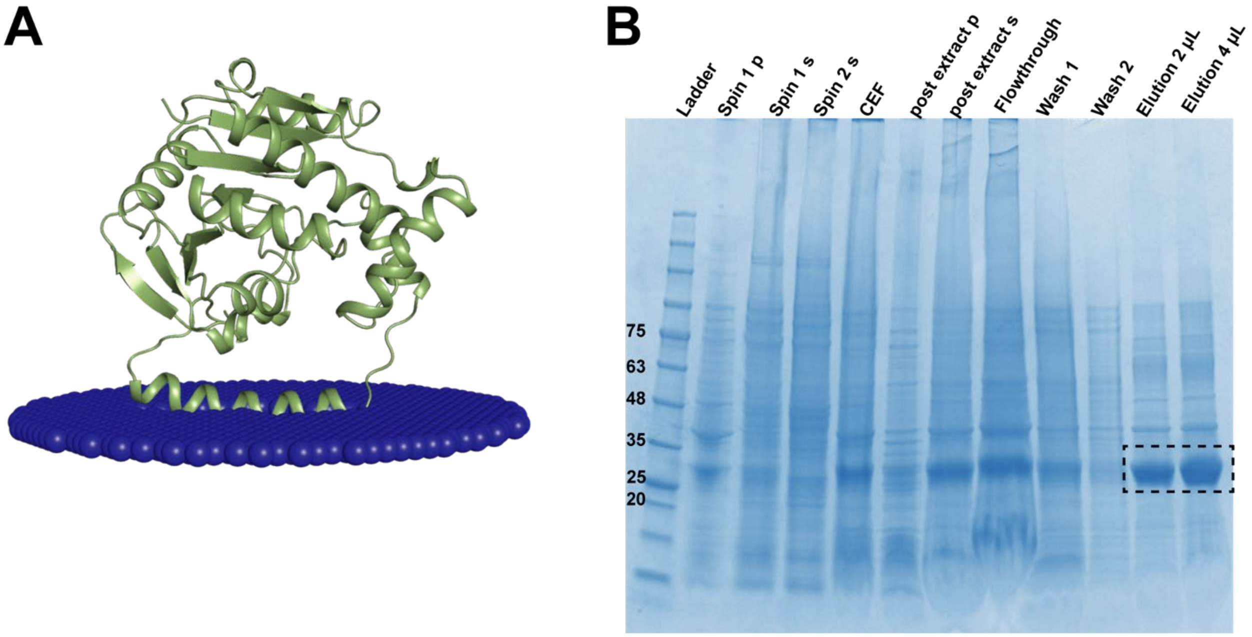


**Figure S2**. GT-25 predicted structure and purification. (A) AlphaFold model of GT-25 (green) docked into the membrane (represented by blue spheres) by the PPM server. (B) SDS-PAGE of GT-25 purification steps. Pellet (p) and supernatant (s) are abbreviated. Volume listed represents the amount loaded.

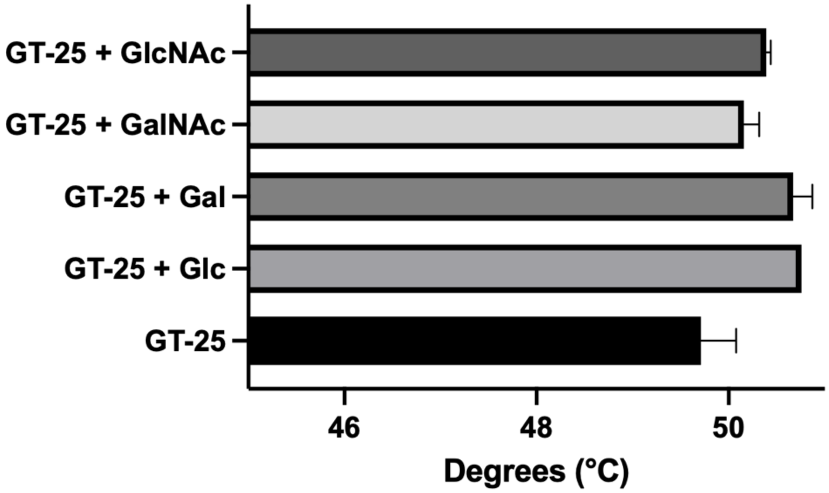


**Figure S3**. NanoDSF of GT-25. Samples were run in duplicate and error bars are shown. For the Glc sample, the error bar is extremely small <0.05 and not shown at this scale.

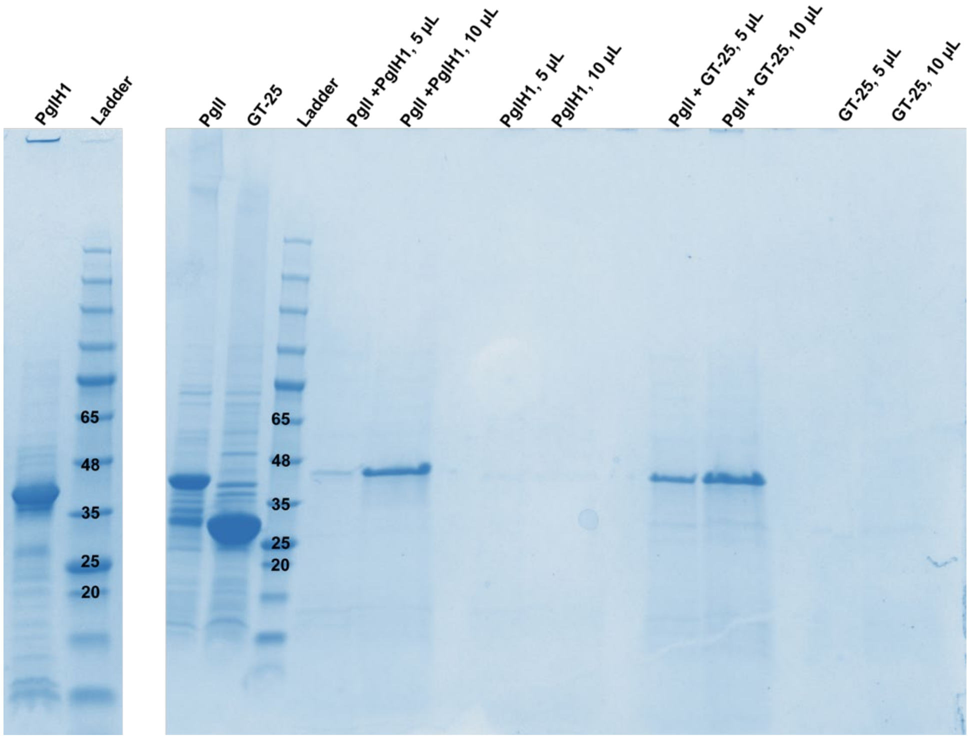


**Figure S4**. Pulldown assay of PglH1, PglI, and GT-25. SDS-PAGE lanes are labeled accordingly. Volumes listed represent the loading volume.

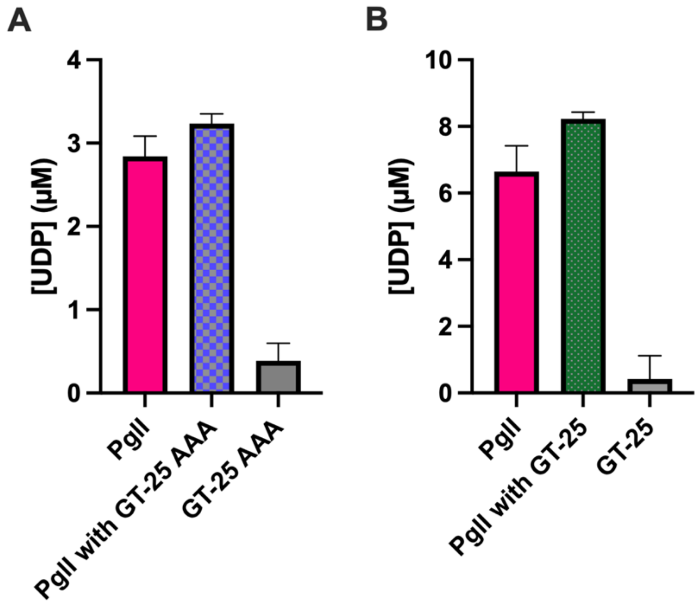


**Figure S5**. UDP-glo activity assays evaluating the effect of GT-25. (A) PglI activity with and without GT-25 AAA variant with GT-25 AAA background activity shown. (B) PglI activity with and without GT-25 with GT-25 background activity shown. Reactions were run in duplicate and error bars are shown.

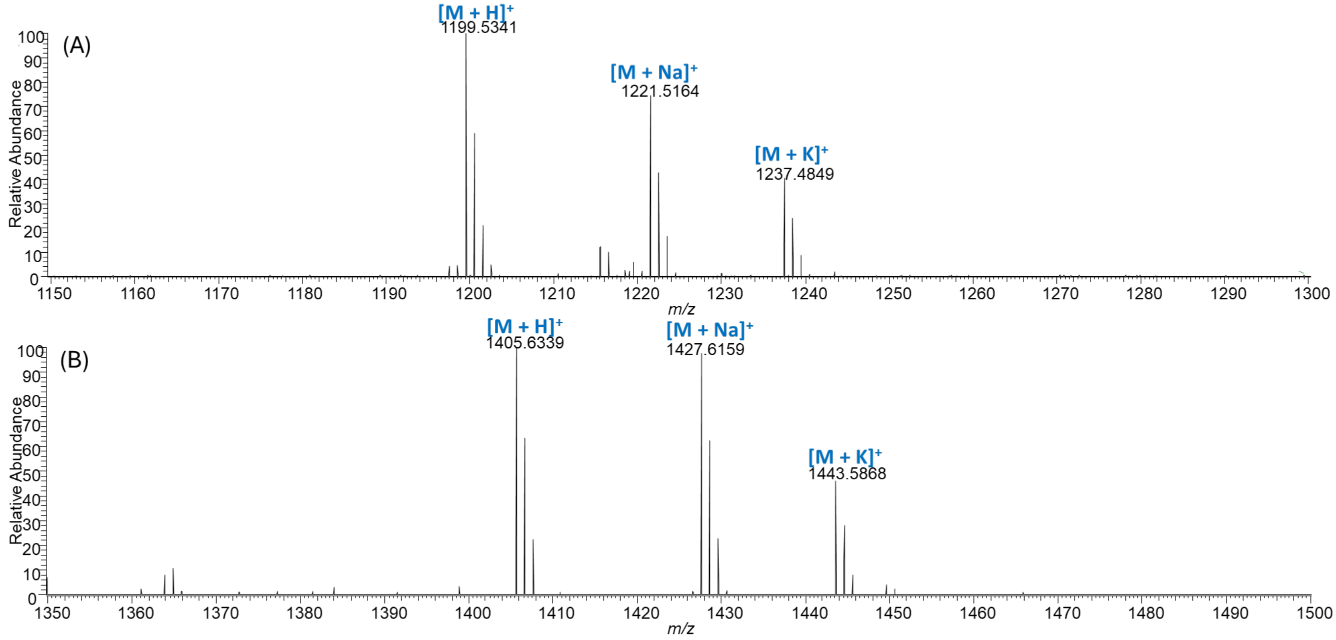


**Figure S6**. Molecular ion region in the positive-ion high resolution MS1 spectra of the products from the (**A**) PglH1 and (**B**) PglI reaction steps.

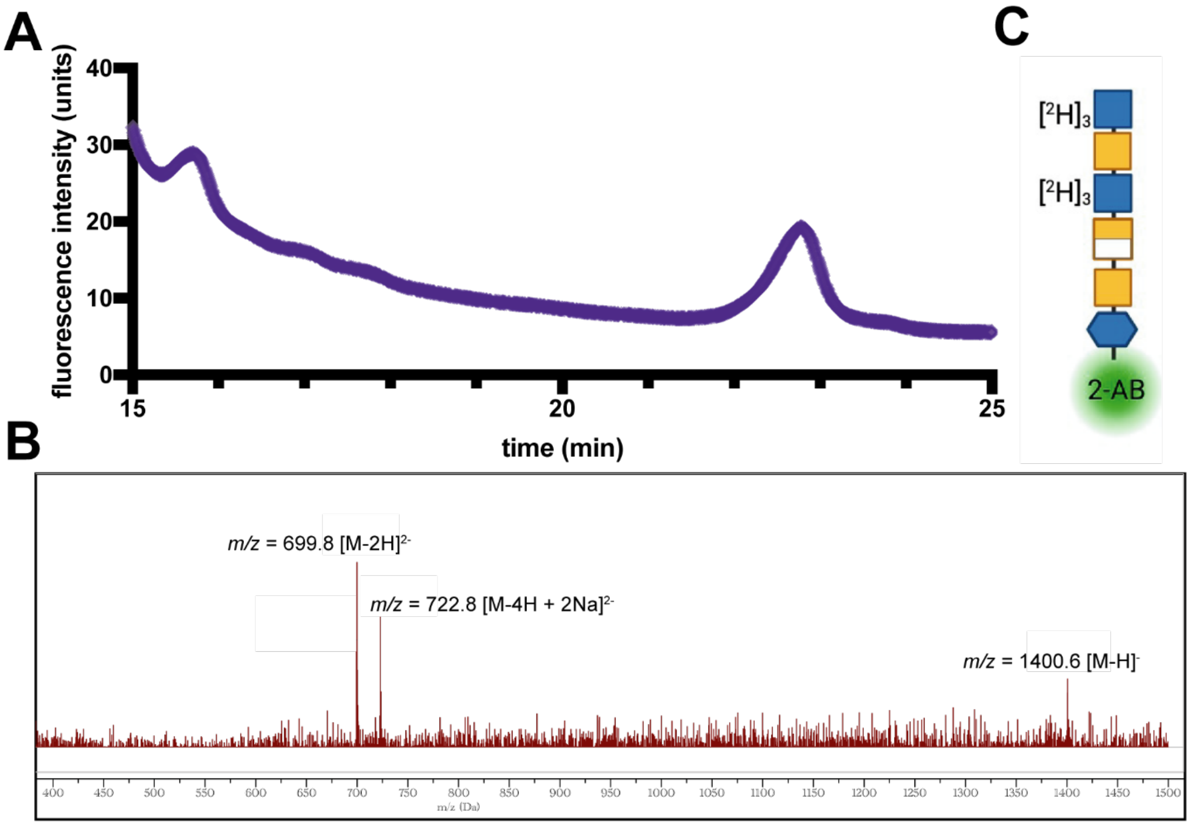


**Figure S7**. Characterization of the product from the chemoenzymatic synthesis with the PglH1 S258P/N259T variant. (A) The fHPLC trace with one peak centered between 22 and 23 minutes. (B) The low-resolution negative-ion mass spectrum [ESI-] obtained for the peak. (C) The glycan diagram showing the structure corresponding to the observed mass.

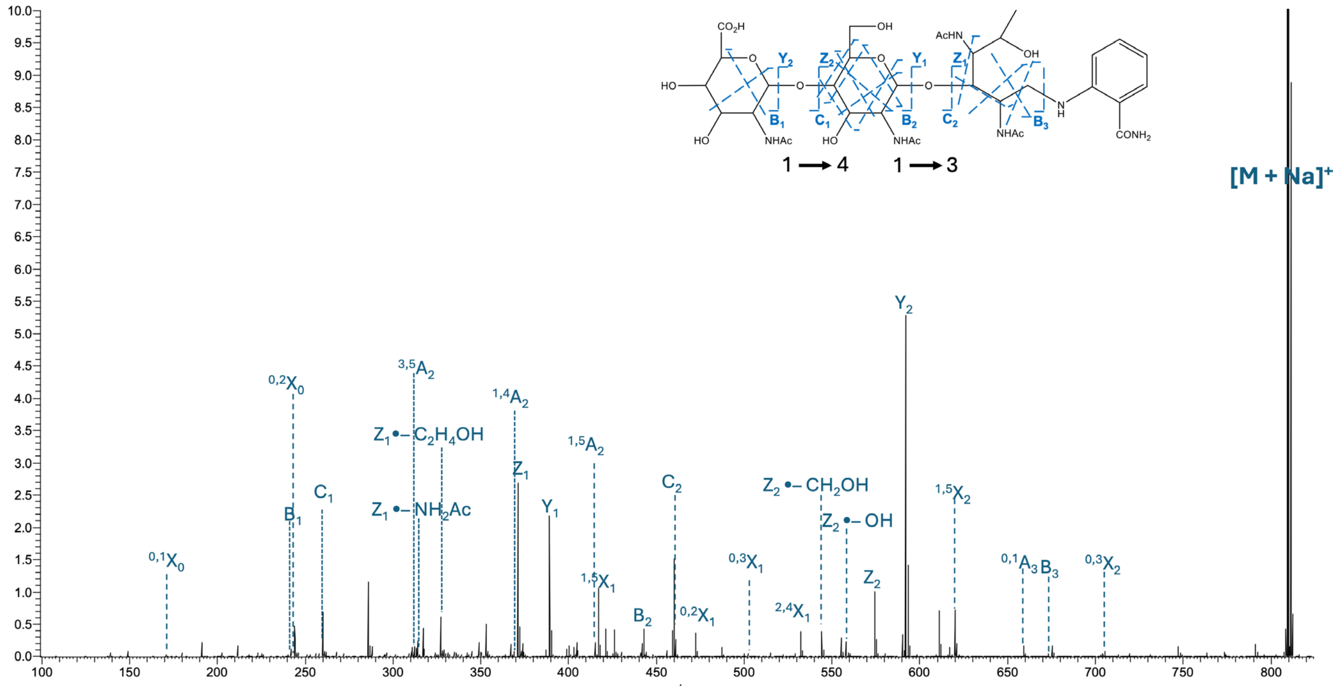

**Figure S8**. EED MS/MS spectrum of the $[M + Na]^{1+}$ *m/z* 809.3156 of the PglJ product.

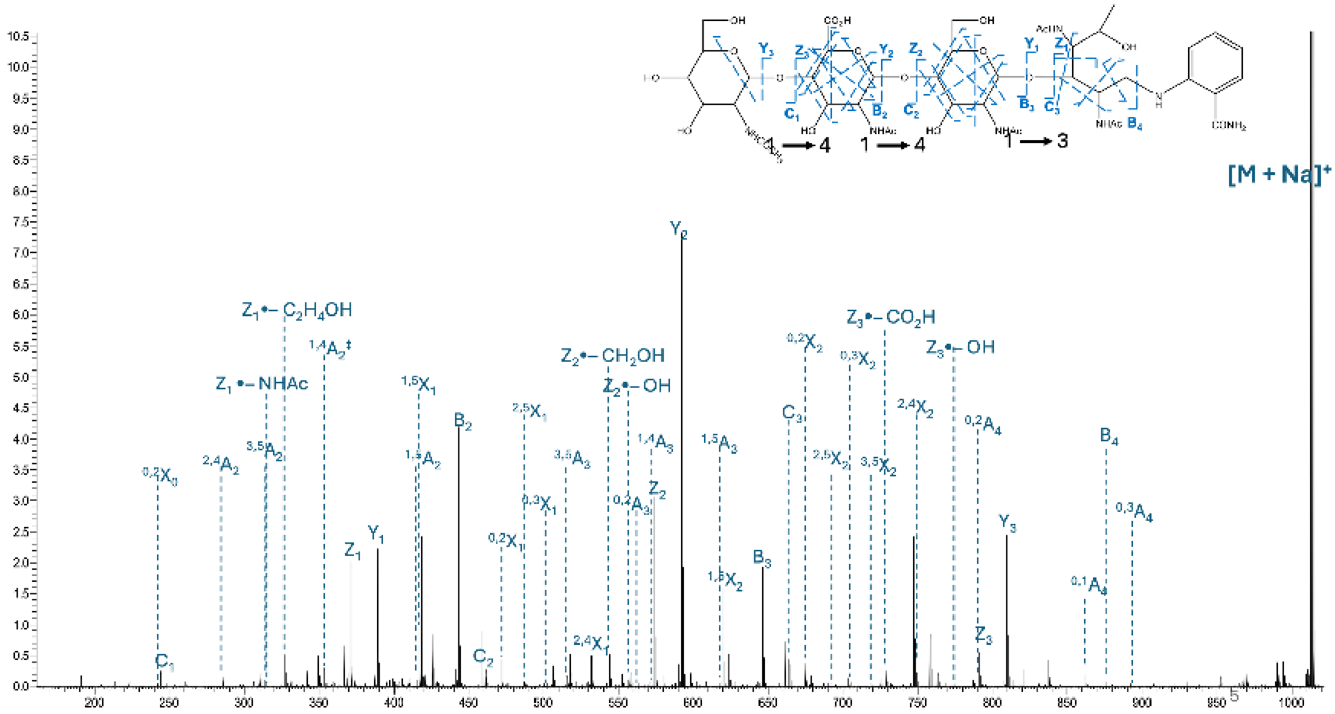

**Figure S9**. EED MS/MS spectrum of the $[M + Na]^{1+}$ *m/z* 1012.3999 of the PglH2 product.

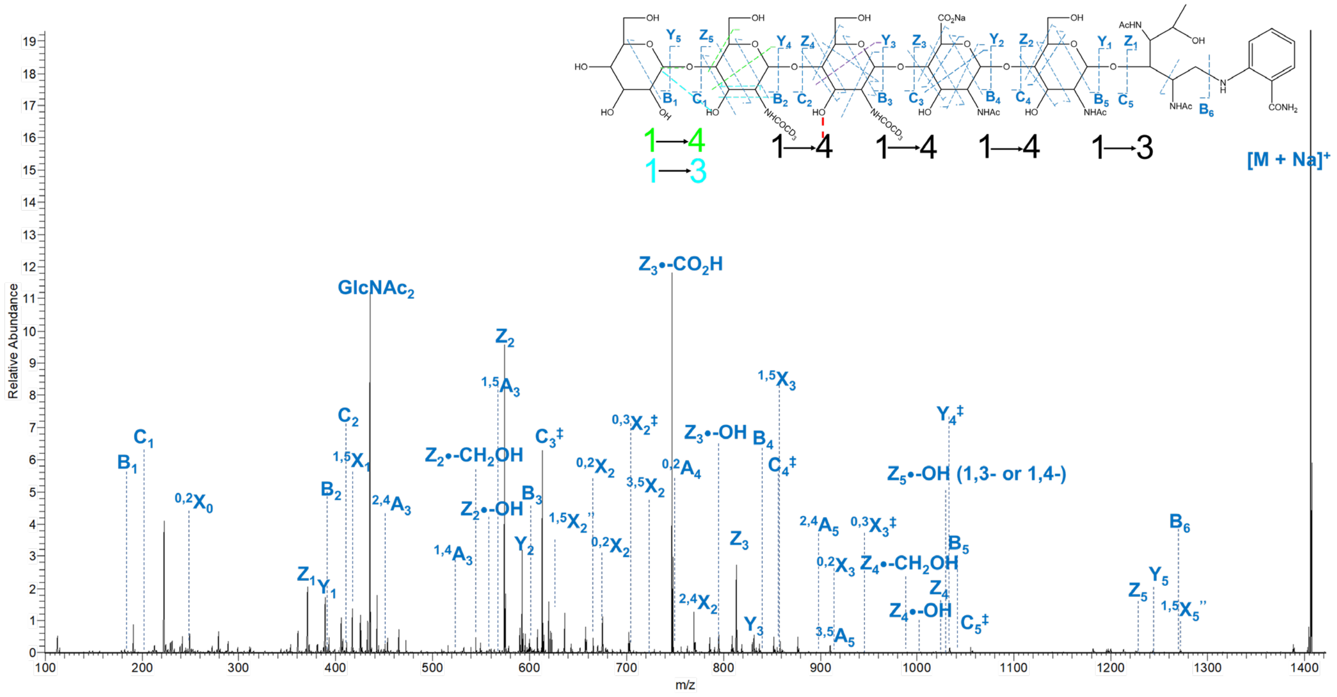


**Figure S10**. EED MS/MS spectrum of the $[M + Na]^{1+}$ *m/z* 1405.5519 of the PglI product with Glc addition.
